# Biochemical, molecular and antibiotic resistance profile of multi-potential toluene metabolizing bacteria isolated from tannery effluents

**DOI:** 10.1101/340240

**Authors:** Fatima Muccee, Samina Ejaz

## Abstract

The focus of present study was to isolate and characterize bacteria which can be effectively used for toluene, a highly recalcitrant pollutant, bioremediation. For isolation of bacteria from the tannery effluents selective enrichment and serial dilution methods were employed. The isolated bacteria were subjected to growth curve analysis, estimation of toluene removal efficiencies, biochemical tests, antibiotic sensitivity assays and molecular characterization based upon 16S rRNA gene. The rRNA genes sequences were analyzed through BLAST to determine similarity index of isolates with bacterial database sequences. To trace the evolutionary history, phylogenetic trees were constructed using MEGA version 7. Total twenty toluene metabolizing bacteria (IUBT1-2, 4-12, 16, 19, 21, 23-26, 28 and 30) were isolated and characterized. Their rRNA gene sequences have been submitted to Genbank. Fifteen of the twenty isolates showed homology to *Brevibacillus agri* strain NBRC 15538, four found similar to *Bacillus paralicheniformis* strain KJ-16 and one homologous to *Burkholderia lata* strain 383. All bacterial isolates were resistant to chloramphenicol but sensitive to teicoplanin and linezolid. However, few (i. e.; IUBT9 and 26) were sensitive to oxacillin. Biochemical characterization indicated all bacteria positive for alkaline phosphatases (100%). While many were found positive for p-nitrophenyl N-acetyl β, D-glucosaminidase (35%), hydroxyproline β-naphthylaminopeptidase (15%), esculinase (65%), mannitol (75%), sorbitol (95%) and inulin (90%) fermentation. Biochemical profile suggests the use of isolated bacteria for future exploitation in several fields like bioremediation of toluene, ethanol production, biomass hydrolysis, biosensors, biofertilizers, as a marker for milk pasteurization in dairy industries and evaluation of soil quality.

**Importance:** Toluene is a highly toxic environmental pollutant. We have isolated bacteria which can be effectively used for the removal of toluene from environmental resources. Moreover, these bacteria are capable to produce many valuable enzymes which can be used in many industrial processes for the production of a wide range of products. Further study may help to exploit these bacterial for the benefit of humanity.

## Introduction

Toluene (methylbenzene) is an omnipresent pollutant and is a cause of concern due to its resistance to chemical, photolytic and biological degradation, lipophilic nature, bioaccumulation, long-range transport and wide range adverse effects on environment, wild life, biota and human health (1). Being lipophilic, it gets accumulated in lipid bilayer of cell membrane and alters the structure of living cells (2). Toluene is a man-made hydrocarbon and its major sources in urban air environment are industrial effluents and motor vehicle emissions (3). It is used as solvent and is essential component of printing inks, adhesives, rubber, disinfectants, leather tanners, petrol, paints, lacquers and cleaning agents. Moreover, toluene is added to gasoline fuels to boost octane number (4).

Breathing in indoor or ambient air around toluene sources is the cause of toluene exposure and may lead to various health issues. Low level exposure (200 ppm) promotes headache, fatigue, slowed reflexes and paresthesia while exposure to moderate levels of toluene (600 ppm) elevate feelings of confusion and high levels (800 ppm) of toluene lead to euphoria (5). The other chronic consequences of toluene exposure include teratogenic effects, neurotoxic and immunotoxic effects, dementia and toluene leukoencephalopathy (6–8).

Due to associated health risks, it is essential to mitigate toluene. Various traditional techniques like soil vapor extraction, incineration, chemical oxidation, ultraviolet oxidation and nanotech remedial technology etc. have been used for this purpose (9–12). However, conventional approaches due to associated limitations like partial toluene degradation, high operating and maintenance cost, maximum disturbance to land, production of more hazardous substances, transport of contaminated material and increased exposure to toluene for the workers are not preferentially used.

The bioremediation has become a preferable technique for toluene detoxification. The process of bioremediation involves scavenging of environmental (water or soil) organic pollutants by naturally occurring or deliberately introduced cultured microbes which are capable to consume and metabolize the contaminants and thus neutralize the environment (13). The bioremedial approach is; far less disruptive and invasive, can be improved by using genetically engineered microorganisms, avoids further pollution, more accessible and cheaper alternative (14). Utilization of bacteria to eradicate environmental pollution is beneficial due to their rapid growth rate, broad metabolic capabilities and diverse enzymes which are active in aerobic as well as anaerobic environments. Moreover bacteria can survive under wide range of temperature and salinity conditions.

Variety of toluene metabolizing bacterial species isolated from different sources has been reported (Table 1).

**Table 1:** Toluene metabolizing bacteria

| Bacterial species | Source | Reference |
| --- | --- | --- |
| *Bacillus cereus* THH39 | Oil polluted soils | (43) |
| *Magnetospirillum* sp. strain 15-1 | Rhizospheres of constructed wetlands. | (44) |
| *Bacterium* Ex-DG74 | Wastewater | (17) |
| *Pseudomonas aeruginosa* UKMP-14T and *Bacillus cereus* UKMP-6G | oil contaminated soil | (45) |
| *Acinetobacter* *genospecies* Tol 5 | gasoline contaminated sediment at a gas station | (46) |
| *Pseudomonas aeruginosa* AT18 | crude oil contaminated soil | (47) |

Being a highly reduced molecule, toluene needs to be oxidized before its assimilation. For this purpose, bacteria have adopted diverse degradative pathways selectively operating under aerobic or anaerobic conditions (Table 2).

**Table 2:** Pathways for aerobic oxidation of toluene

| Pathway | Bacteria | Reference |
| --- | --- | --- |
| Dioxygenase mediated pathway | *Thauera* species strain DNT-1 | (48) |
| Toluene-2-monooxygenase pathway (TOM) | *Burkholderia cepacia* G4 | (49) |
| Toluene-3-monooxygenase pathway (TBU) | *Ralstonia pickettii* PKO1 | (50) |
| Toluene-4-monooxygenase pathway (TMO) | *Pseudomonas mendocina* KR1 | (51) |
| Ortho cleavage pathway | *Pseudomona mendocina* KR1 | (27) |
| Meta cleavage pathway via catechol | *Burkholderia cepacia* G4 | (26) |
| Tod pathway | *Pseudomonas putida* F1 | (52) |

Although a number of toluene metabolizing bacteria have been isolated and studied, there is need to find more highly efficient and environment friendly bacteria. Under different habitats, bacteria express different biodegradative enzymes and pathways (15). Moreover, utilization of bacterial consortia with broad spectrum of enzymes will be proven more effective for toluene remediation (16).

The industries which manufacture and use toluene are the primary sources of toluene pollution in the environment. Toluene is a commonly used adhesive in leather industry. Hence, leather tanning and processing industries and manufacturing units of footwears, leather bags, leather furniture and other miscellaneous leather goods are the major sources of toluene exposure. Knowing thing in present study, tannery effluent was selected as a source sample for the isolation of toluene metabolizing bacteria. It was hypothesized that we might find some novel bacteria capable to degrade toluene in a more effective and versatile manner in tannery industry effluents. The accomplishment of this project might provide an eco-friendly way out to reclaim toluene contaminated environmental resources.

## Materials and methods

### Medium and growth conditions

For bacterial isolation, standard culture enrichment technique was employed by involving use of carbon deficient minimal salt media (17). The carbon deficient minimal salt medium (4g L^−1^ KH_2_PO_4_; 4 g L^−1^ Na_2_HPO_4_; 2 g L^−1^ NH_4_Cl; 0.2 g L^−1^ MgCl_2_; 0.001 g L^−1^ CaCl_2_; 0.001 g L^−1^ FeCl_3_) was prepared and autoclaved at 121 °C for 20 min. Prior to inoculation, the medium was supplemented with 1% (v/v) toluene as the only carbon source. The pH of medium was maintained at 6.8. Isolated bacteria were grown aerobically on this toluene supplemented media at 50 °C.

### Morphology

Morphological aspects of isolates like gram staining and bacterial cell shape were documented. The colony forming units (CFUs) were also calculated for all these isolates.

### Growth rate

The growth rate was evaluated indirectly by assessment of turbidity as optical density (OD). The optical density was measured at 600 nm and at different time intervals using UV-visible spectrophotometer. The optical density was plotted versus time to obtain growth curves and determine growth behavior of each isolate.

### Toluene removal assay

For determining the toluene removal rate of bacterial isolates, initially the bacterial culture was centrifuged. The supernatant containing the residual toluene of the medium (3ml) was extracted using 3ml n-hexane. Followed by this, the n-hexane containing toluene was separated using a separating funnel. The optical density of toluene was measured against a blank at 260 nm wavelength (Supplementary Data Figure 1). Medium containing toluene without bacterial culture was used as blank. Toluene degradation efficiency has been calculated by following formula;

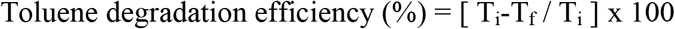

Whereas; T_i_ = initial absorbance of toluene

T_f_ = final absorbance of toluene

### Morphological and Biochemical characterization

The isolated bacteria were characterized by gram staining, colony morphology (18) on nutrient agar medium and morphological characteristics. For biochemical identification of bacteria, a qualitative micromethod of Remel RapID STR System (19) was used. In this method, toluene metabolizing bacteria were subjected to different biochemical tests. i.e.; arginine dehydrolase test, esculinase test, mannitol fermentation test, sorbitol fermentation test, raffinose fermentation test, inulin fermentation test, ρ-nitrophenyl-α,D-galactosidase test, Tyrosine β-naphthylamidase test, ρ-Nitrophenyl-α,D-glucosidase test, ρ-Nitrophenyl-n-acetyl-β,D-glucosaminidase test, Lysine β-naphthylamidase test, alkaline phosphatase test and pyrrolidonyl peptidase test.

### Antibiotic susceptibility test

Antibiotic test was performed for all the isolated bacteria by using disc diffusion method (20). Antibiotics tested were teicoplanin (30 μg), linezolid (30 μg), linezolid (10 μg), oxacillin (1 μg) and chloramphenicol (30 μg).

### Molecular characterization by 16S rRNA gene sequence analysis

For molecular characterization of bacterial isolates their genomic DNA was extracted by organic method (21). The 16S rDNA gene was amplified by polymerase chain reaction (PCR) using the rDNA specific primers F1 and R1 (Supplementary Data Table 1). The region of 16S rRNA gene targeted by these primers is shown (Supplementary data Figure 2). In 50 μl of PCR reaction mixture, 50 ng of template DNA, 5 ul of 10X PCR reaction buffer (Mg^++^ free), 5 μl of MgCl_2_, 1 μl of 10mM dNTPs, 2 μl of 10 pM forward primer, 2 μl of 10 pM reverse primer, 0.25 μl of Taq DNA polymerase and 29.75 μl of nuclease free water was used. PCR amplification conditions were as follows; initial denaturation at 95 °C for 5 min, 38 cycles consisting of denaturation at 94 °C for 40 seconds, annealing at 58 °C for 40 seconds and extension at 72 °C for 30 seconds. The final extension of PCR products was carried out at 72 °C for 10 min. After PCR amplification, the PCR products were resolved on agarose gel. The PCR products were purified by Monarch DNA Gel Extraction Kit and sequenced bidirectionally using sequencer based on principle of Sangers Chain Termination. For this purpose, samples were sent to Macrogen, Korea. FASTA sequences obtained from Macrogen were subjected to microbial databases similarity searches through NCBI (National Center for Biotechnology Information) BLAST analysis (http://blast.ncbi.nlm.nih.gov/blast/Blast.cgi). Sequences of highest similarity obtained through BLAST analysis were recorded. Clustal omega multiple sequence alignment software (22) was used to align the sequences. The sequences were refined by removing the gaps and only ungapped aligned regions were analyzed by MEGA7 software to perform phylogenetic analysis of the identified bacteria. Phylogenetic trees were constructed to show the evolutionary relationship among the identified bacteria. Different parameters used for phylogenetic tree construction included neighbor joining statistical method, maximum composite likelihood substitution method and Bootstrap analysis with 1000 replicates.

Sequences of twenty isolates have been submitted to NCBI and accession numbers assigned are; IUBT1 (MH014864), IUBT2 (MG241284), IUBT4 (MH014871), IUBT5 (MH014859), IUBT6 (MH266116), IUBT7 (MH201270), IUBT8 (MG190354), IUBT9 (MG190357), IUBT10 (MG263995), IUBT11 (MH266117), IUBT12 (MG263996), IUBT16 (MG190391), IUBT19 (MH014861), IUBT21 (MH014862), IUBT23 (MG263997), IUBT24 (MG190870), IUBT25 (MH266118), IUBT26 (MG263998), IUBT28 (MH014868) and IUBT30 (MH014869).

## Results

### Morphology

All the bacteria were found to be gram positive and rod shaped. Their Colony Forming Units (CFUs) have been calculated and are given (Supplementary Data Table 2). The bacteria IUBT1, IUBT2, IUBT4 and IUBT5 were isolated from the plate inoculated with original culture without any dilution.

### Molecular characterization of toluene metabolizing bacteria

Ribotyping of toluene metabolizing bacteria revealed three groups among twenty isolates (Figure 1). Group 1 comprised of single isolate IUBT1, homologous to *Burkholderia lata* strain 383. Group 2 included fifteen bacteria homologous to *Brevibacillus agri* strain NBRC 15538 while group 3 contained isolates IUBT4, IUBT24, IUBT28 and IUBT30 showing homology to *Bacillus paralicheniformis* strain KJ-16. All sequences were deposited to Genbank database (www.ncbi.nlm.nih.gov/genbank) and assigned accession numbers.

**Figure 1:**
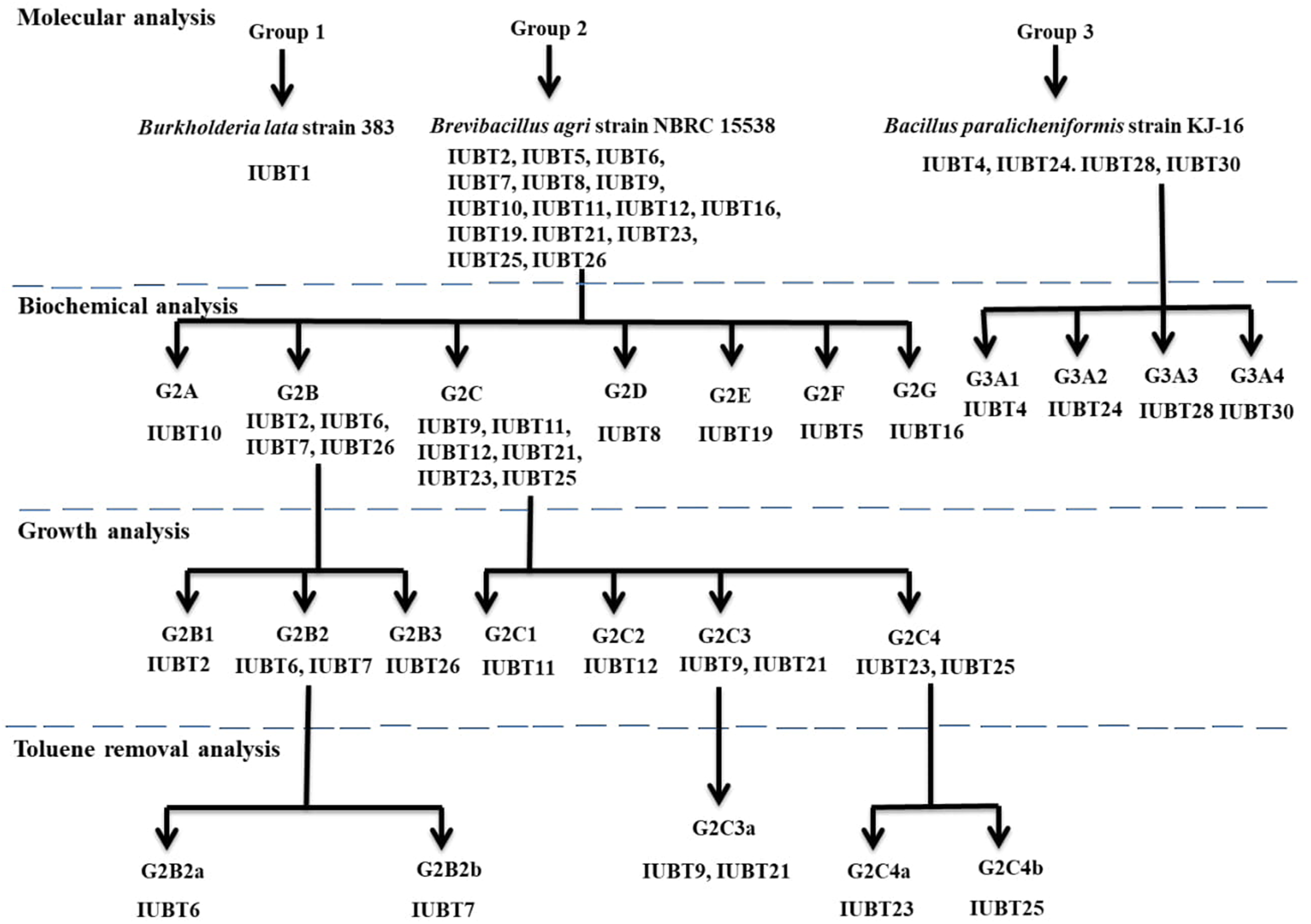
Flowsheet for twenty toluene metabolizing bacteria: The molecular, biochemical, growth pattern and toluene removal analysis data showed eighteen bacteria IUBT1, IUBT2, IUBT4, IUBT5, IUBT6, IUBT7, IUBT8, IUBT9, IUBT10, IUBT11, IUBT12, IUBT16, IUBT23, IUBT24, IUBT25, IUBT26, IUBT28, IUBT30 were unique. And two bacteria IUBT9 and IUBT21 might be the same on the basis of similarity in their toluene removal efficiencies.

### Phylogenetic analysis of bacterial isolates

To study the evolutionary relationships between the isolated toluene metabolizing bacteria phylogenetic trees were constructed. Bacterium IUBT1 from group 1, was assigned to genus *Burkholderia*. Study of its evolutionary relationship revealed its common ancestry with *Burkholderia lata strain 383* (Accession number NR 102890) as both originated from the same node (Figure 2). The similarity percentage between two strains was 99%.

**Figure 2:**
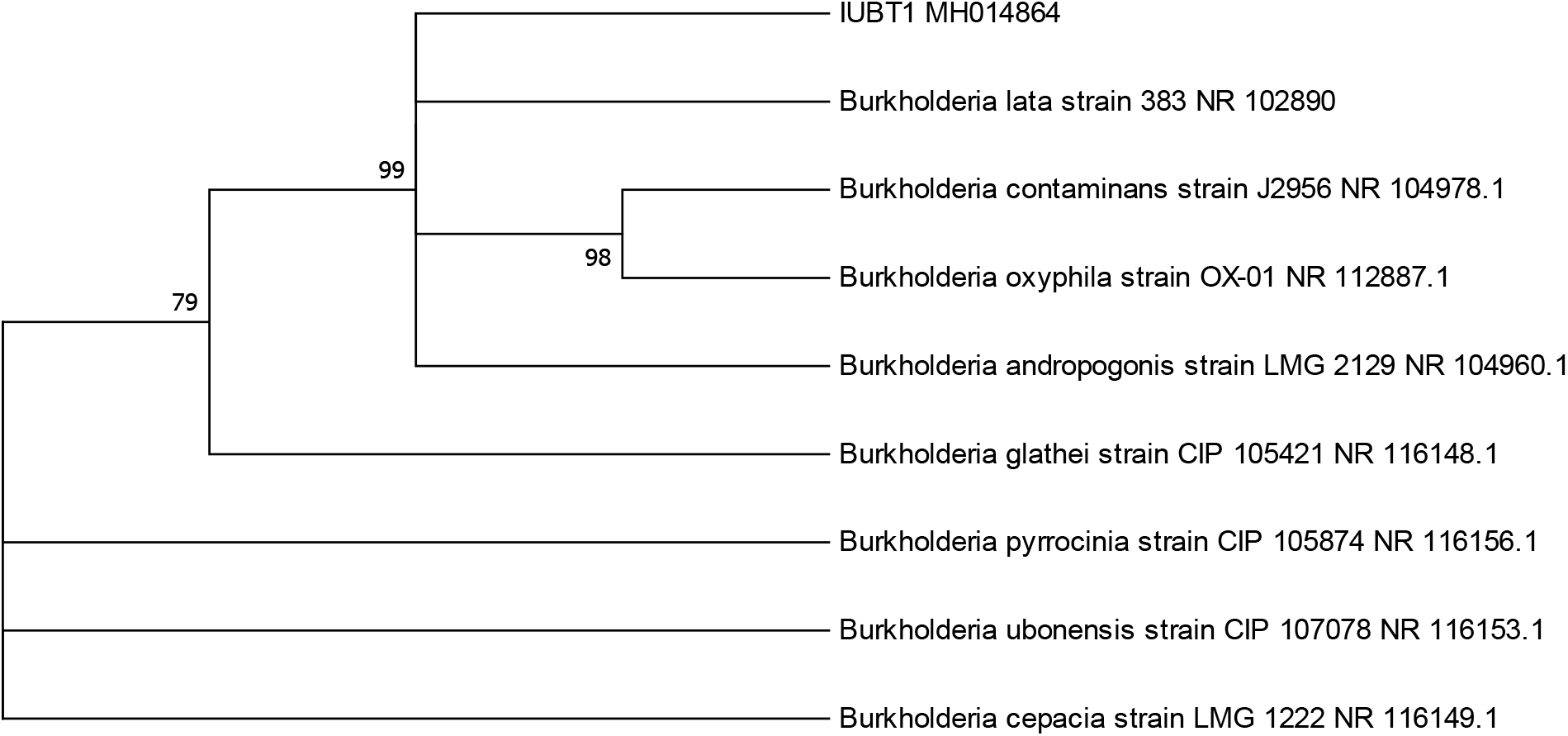
Neighbour joining tree based on 16S rRNA gene sequence for IUBT1 and its closest relatives. Numbers with the tree nodes are depicting percentage bootstrap support for 1000 replicates.

Four bacterial isolates IUBT4, IUBT24, IUBT28 and IUBT30 were assigned to genus *Bacillus*. All shared common ancestor, belonged to same clade and shared close similarity with *Bacillusparalicheniformis* strain KJ-16 (Figure 3).

**Figure 3:**
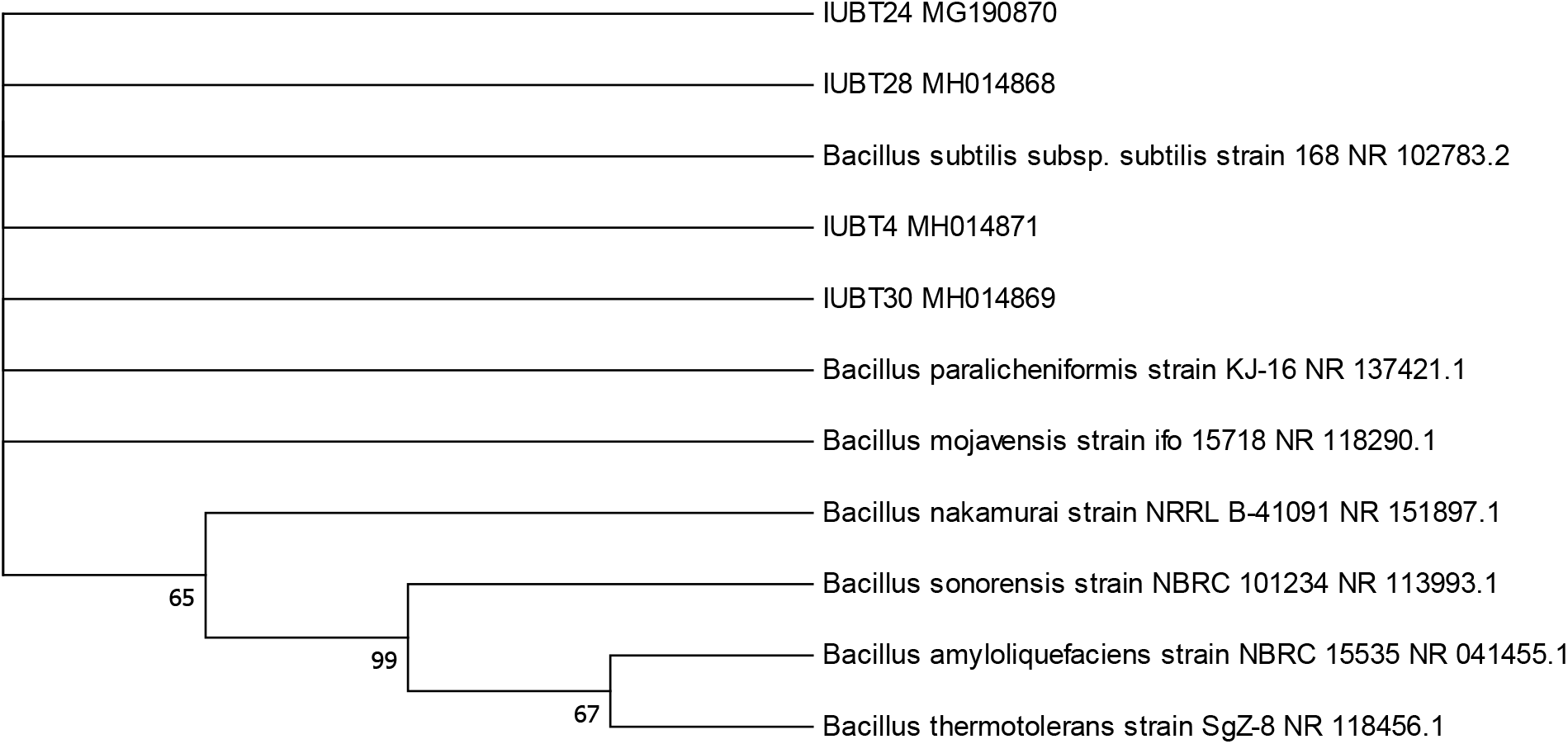
Neighbour joining tree illustrating the evolutionary relationship between IUBT4, IUBT24, IUBT28, IUBT30 and other related strains.

Among the 20 isolates in this study, 15 were assigned to genus *Brevibacillus* in Group 2. The phylogenetic study revealed that IUBT7 is distantly related with rest of the isolates and same is the case of IUBT9. Both of these are not sharing clade with any other isolate. IUBT12 and 19 are sharing the same clade. IUBT21 and IUBT23 are related through strong bootstrap value of 99. IUBT5 and IUBT11 are also closely related because they are converging at a common ancestor. IUBT6 is distantly related with IUBT26 through bootstrap value of 22. IUBT2 is related with *Bacillus glycinifermentans* through strong bootstrap value of 100. IUBT8 and IUBT10 are more related with each other as compared to IUBT16 and IUBT25. IUBT25 is sharing the same lineage with *Brevibacillus gelatini* strain through bootstrap value of 48 (Figure 4).

**Figure 4:**
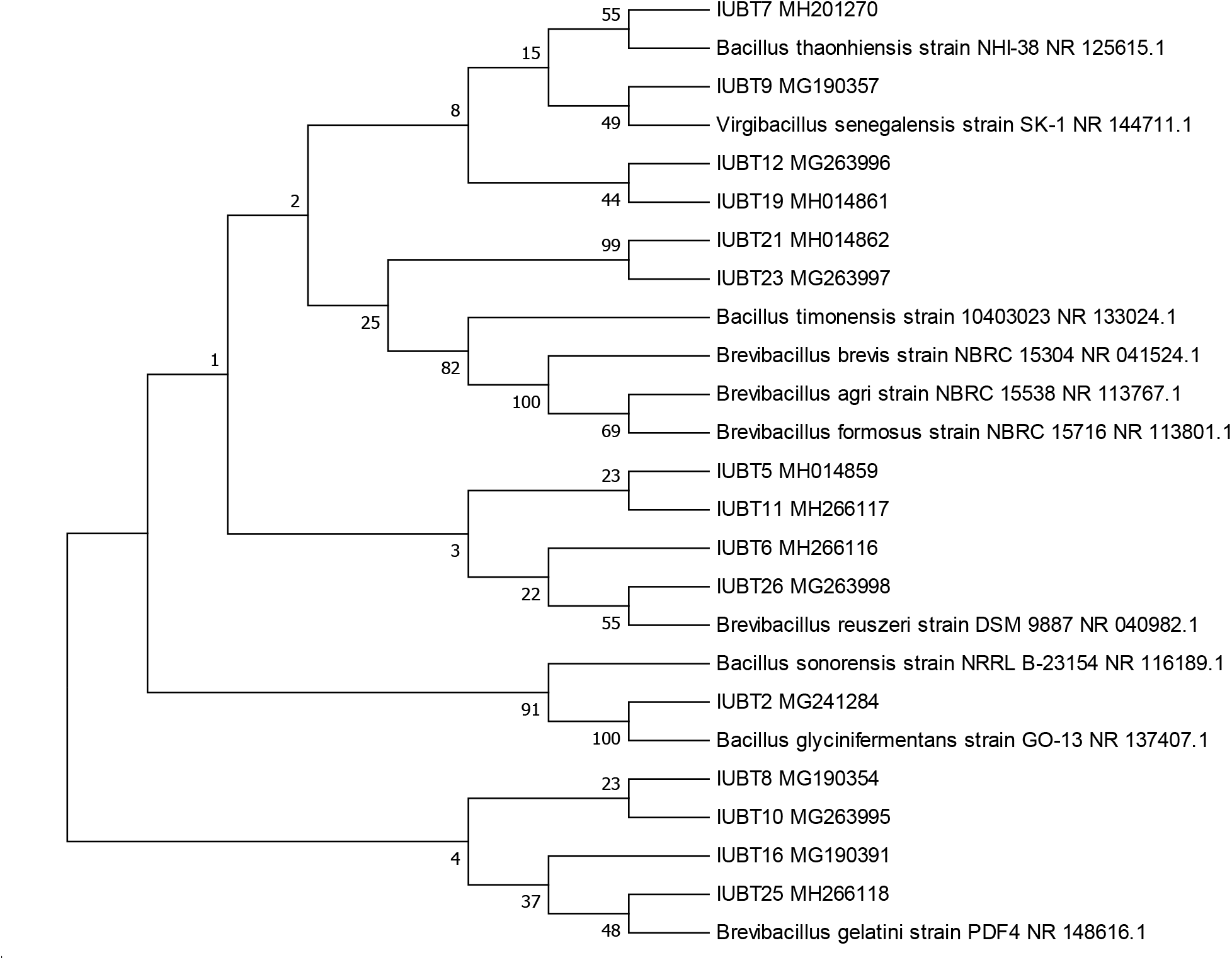
Neighbour joining tree illustrating phylogenetic relationship of isolates assigned to genus *Brevibacillus* and other related strains.

### Biochemical characterization

The isolated bacteria were screened for fourteen biochemical tests. All of these were negative for arginine dehydrolase (ARG) test, raffinose fermentation (RAF) test, ρ-nitrophenyl-α, D-galactoside (GAL) test, tyrosine β-naphthylamide (TYR) test, ρ-nitrophenyl-α, D-glucoside (GLU) test and lysine β-naphthylamide (LYS) test. The positive response was observed for the esculinase (ESC) test, mannitol fermentation test (MNL), sorbitol fermentation test (SBL), inulin fermentation test (INU), hydroxyproline β-naphthylamide (HPR) test, ρ-nitrophenyl-n-acetyl-β-D-glucosaminide (NAG) test, ρ-nitrophenyl phosphate (PO_4_) test and pyrrolidine β-naphthylamide (PYR) test. Thirteen bacteria were esculinase producers and can be exploited for improved ethanol production through increased saccharification of cellulose. All except two (IUBT8 and 16) were capable of fermenting inulin and fifteen (IUBT2, 4, 5-7, 9, 11-12, 19, 21, 2-26 and 30) were mannitol fermenting bacteria. All the isolates were positive for sorbitol fermentation except IUBT16. Seven were n-acetyl glucosaminidase producers and possess potential for future applications like single cell protein (SCP) production, for conversion of chitinous waste into biofertilizers and as an additive in food industry. Only two (IUBT5 and 19) were hydroxyproline β-naphthylamidases producers and can be utilized in pharmaceutical industeries. All isolates were positive for PO4 test and thus can be employed in diagnostics immunoassays, in biosensors for environment monitoring, as biofertilizers, in soil quality evaluation, in limnological studies and dairy industries. Only one bacterium IUBT5 was positive for PYR test and can be utilized in protein sequencing to free the amino terminal amino acid being blocked by pyrrolidonyl group. This analysis helped us to find out unique characteristics of Group 2 and Group 3 members. Biochemical behavior of IUBT10, IUBT8, IUBT19, IUBT5 and IUBT16 was different from each other and from ten other *Brevibacilli* of Group 2. So, group 2 members were further categorized into two major groups. The isolates IUBT2, IUBT6, IUBT7 and IUBT26 were similar biochemically so placed in the group G2B. These bacteria were sorbitol and inulin fermenting and esculinase, p-nitophenyl-n-acetyl-β-D-glucosaminidase and p-nitrophenyl phosphatase producers. The IUBT9, IUBT11, IUBT12, IUBT21, IUBT23 and IUBT25 were placed in group G2C as they all were mannitol, sorbitol and inulin fermenting and esculinase and p-nitrophenyl phosphatase producers. All the four isolates from Group III were found biochemically different. Hence, biochemical analysis confirmed their distinctiveness (Supplementary Data Table 3).

### Growth study

To sort out the unique bacteria among those having same molecular and biochemical profile their growth behavior was studied. Assuming that same bacteria must possess same growth pattern, the isolates IUBT2, IUBT26, IUBT11 and IUBT12 were found unique in their growth behavior (Supplementary data Table 4). Results helped to identify three groups of bacteria G2B2 consisting of IUBT6 and IUBT7, G2C3 containing IUBT9 and IUBT21 and G2C4 including IUBT23 and IUBT25 (Supplementary Data Figure 3). The members of each group exhibited similar growth pattern hence, they might be the same bacteria.

### Toluene removal efficiency

Toluene removal efficiency assay served as fourth layer of screening and helped to differentiate among bacteria exhibiting similarity in molecular, biochemical and growth behaviors. The toluene removal potentials for IUBT6, IUBT7, IUBT23 and IUBT25 were found to be 83% ± 1.41, 93% ± 4.95, 28% ± 4.24 and 45% ± 10.6 respectively (Supplementary data Figure 4). The IUBT9 and IUBT21 had toluene removal potentials: 83% ± 2.12 and 83% ± 0.70, respectively. However, IUBT9 and IUBT2 might be the same bacteria due to similarity in their molecular aspects, biochemical behavior, growth pattern and toluene removal efficiencies.

### Antibiotic profile

All of the twenty isolates were tested for their antibiotic susceptibility (Supplementary Data Table 5). All of the isolates were sensitive to teicoplanin and linezolid. Among these isolates, 100% were resistant to chloramphenicol and 90% to oxacillin. Only two of these, IUBT9 and IUBT26 showed sensitivity to oxacillin and 100% showed sensitivity to teicoplanin and linezolid. The zone of inhibition was measured for the antibiotics to which bacteria exhibited susceptibility. The maximum zone of inhibition was recorded 39 mm against the antibiotic linezolid (10μg) in case of IUBT19. In case of teicoplanin, linezolid (30μg) and oxacillin, maximum zones of inhibition were 35 mm (IUBT6), 37 mm (IUBT16, 26 and 30) and 30 mm (IUBT26), respectively. A comparison of zone of inhibitions for different antibiotics is given (Supplementary Data Figure 5).

## Discussion

Grown on minimal salt medium (MSM) supplemented with 1% toluene, morphological, biochemical and molecular analyses of all isolates (n=20) revealed their capability to metabolize toluene as the only source of carbon.

### Morphology

Morphologically these strains were found to be rod shaped and gram positive bacteria. Our results are consistent with earlier studies because most of the toluene metabolizing bacteria reported in literature are gram positive: Bacterium Ex-DG74 (17), *Mycobacterium sp*. IBB_pol_ (23), *Mycobacterium* strains T103 and T104 (24). However, contrary to our findings previous studies have reported few toluene metabolizing gram negative bacteria (23).

### Molecular characterization

Traditional bacterial identification based on phenotypic characteristics is not a precise method in comparison with genotypic tools. Similarly through the 16S rRNA gene profiling the poorly described, phenotypically anomalous, rarely isolated strains and even uncultured strains can be identified (25). In present study, the molecular characterization of isolates showed homology with three different genera and species. i. e.; *Brevibacillus agri, Burkholderia lata* and *Bacillus licheniformis*. The *Burkholderia* and *Bacillus licheniformis*, are known to possess toluene degradation ability (26–28). Although a thermophilic bacterium (growth temp. 45 °C) *Brevibacillus agri* strain 13 has been reported to grow and tolerate high concentrations of toluene (5% and 20%, v/v) but it was not capable of utilizing toluene as sole carbon source (29). Hence, to the best of our knowledge, this is the first study reporting isolation of toluene metabolizing *Brevibacillus agri* strain. Similarity score of less than 97% could be considered as an indicative of a new species within a known genus. Moreover, isolate IUBT7 with 89% similarity could be considered as an archetype of new species within the genus *Brevibacillus*.

### Biochemical characterization

To better comprehend the ongoing metabolic activities isolates were subjected to biochemical analysis. Although previous studies have performed biochemical characterization of toluene metabolizing bacteria through standard biochemical tests (30, 31). Esculinase, hydroxyproline β-naphthylamidase and alkaline phosphatase activities of toluene degrading bacteria, have not been yet studied.

Esculinase (ESC) test is an assay used for the detection of beta-glucosidase enzyme, also known as esculinase. It hydrolyzes esculin into esculetin and dextrose. Followed by this esculetin and ferric chloride of the medium react and black brown color is produced. Thirteen isolates (IUBT1-2, 6-7, 9-12, 16, 21, 23, 25-26) showed positive result for esculinase. Few gram positive strains like *Enterococcus faecium, Enterococcus faecalis* and *Streptococcus bovis* are known to be positive for esculinase test. This test aids in differentiating among few families of *Enterobacteriaceae*. Esculinase or β-glucosidase enzyme due to its hydrolytic activity can be effectively used for biofuel production, synthesis of antitumor agent (aglycone moiety), for removing bitterness from citrus fruit juices, cooked soybean syrup, unripe olive and for detoxification of cassava. By reverse hydrolysis, this enzyme can also be used for synthesis of o-alkyl glucoside which can be used in food industry, extraction of organic dyes, in cosmetics and pesticides formulation (32). In this study, thirteen isolates (IUBT1-2, 6-7, 9-12, 16, 21, 23 and 25) have been found positive for this test. As esculinase enzyme has wide range of applications, hence the isolates found positive hold multipurpose potential.

Mannitol, sorbitol, raffinose and inulin fermentation tests were performed to detect the capability of bacteria to utilize the carbohydrate substrates like mannitol, sorbitol, raffinose and inulin respectively. Bacteria usually transform mannitol, reffinose and inulin into acidic products which reduce the pH of medium, change the color of indicator added to the medium. Our 75% isolates were positive for mannitol test, 95% were positive for sorbitol test, 90% were positive for inulin test while none of the isolates showed positive result for raffinose test. In literature, toluene metabolizing bacteria have been reported to exhibit mannitol and sorbitol fermenting potential (31). However, this is the first study reporting on inulin and raffinose fermenting capability of toluene metabolizing bacteria. Bacterial inulases have tremendous industrial applications and can be utilized for the production of fructose syrup, gluconic acid, inulooligosaccharides, lactic acid, mannitol, ethanol and 2,3-butanediol (33). Hence, the isolates found to possess inulases can be exploited for these purposes.

Hydroxyproline β-naphthylamide (HPR) test detects the presence of hydroxyproline β-naphthylamidase enzyme which catalyzes the hydrolysis of hydroxyproline β-naphthylamide and β-naphthylamine. In present study, three bacterial isolates were found positive for this test.

The ρ-Nitrophenyl-n-acetyl-β-D-glucosaminide (NAG) test is a biochemical assay to detect ρ-Nitrophenyl-n-acetyl-β-D-glucosaminidase (NAGase) in bacteria. This enzyme hydrolyzes p-nitrophenyl substituted glycoside and releases ρ-Nitrophenol. In present study, seven isolates showed positive result for this test. Few toluene metabolizing and glucoaminidase producing strains have been reported (31). A strain *Burkholderia pseudomallei* is also known for the production of this enzyme (34). This enzyme can be employed for analyzing complex sugar chains in glycopeptides and glycoproteins, synthesis of variety of biologically important compounds and as biocontrol agents etc (35).

The ρ-nitrophenyl phosphate (PO4) test confirms the presence of phosphatase enzyme which expedites the breakdown of ρ-nitrophenyl phosphate to yellowish ρ-nitrophenol. All the bacterial isolates of our study analyzed for this test showed positive result. This enzyme has diverse applications such as in immunoassays, biomedical industry due to its resistance to denaturation, biosensors for environmental monitoring, as a marker for adequate milk pasteurization, bioremediation agent to mineralize organophosphates and evaluation of heavy metals precipitation from industrial effluents (36)

Pyrrolidine beta-naphthylamide (PYR) test facilitates the detection of enzyme L-pyrroglutamyl amino-peptidase found in *Streptococcus pyogenes* which hydrolyzes the pyrrolidine β-naphthylamide yielding β-naphthylamide. In present study, only one bacterium showed positive result for this test. This enzyme has potential applications in protein sequencing to release the amino acid blocked by pyrrolidonyl group at N-terminus and bacterial diagnosis (37).

The isolates of present study being the source of variety of enzymes can be destined for utilization in scientific and industrial entities. They can be employed in areas like clinical diagnosis, food industry, dairy technology, agroecosystem, molecular biology, genetic engineering and environmental remediation of pollutants.

### Growth study

Majority of the isolates (n=14) were slow growing and found to exhibit logarithmic growth until 48 hours (IUBT3, 19 and 24), 51 hours (IUBT1, 26 and 28), 68 hours (IUBT5, 8-9, 12, 14 and 21) and even until 74 hours (IUBT30). Consistent with our study, four slow growing toluene metabolizing bacteria, exhibiting slow growth rate with exponential growth phase until 90 hours, have been reported (38). The isolates IUBT2, IUBT4, IUBT5, IUBT6 and IUBT8 showed 0.4-0.5 optical density (OD) at 600 nm during the log phase which is consistent with the optimum optical density (OD) of toluene metabolizing bacteria reported in literature (38). The maximum OD at 600 nm reported for toluene metabolizing bacteria so far is 0.8 (17). While in present study IUBT16 and IUBT25 showed OD of 0.9 and IUBT9, IUBT12, IUBT21, IUBT23, IUBT24 and IUBT28 showed OD of 1.0. Hence, present wok reports the maximum biomass growth of toluene metabolizing bacteria.

### Toluene removal efficiency

Toluene removal efficiencies of bacteria were determined by toluene removal assay. The isolates having highest toluene removal efficiency (Figure 2) were IUBT16 (93% ± 1.41), IUBT1 (90% ± 2.83) and IUBT19 (90% ± 0), IUBT2 (89% ± 1.41), and IUBT4 (89% ± 0.71). The observed toluene removal efficiencies are consistent with those reported in earlier literature. i.e.; 92.4% for bacterium J2, 84.8% for bacterium J6 (39). However, the highest toluene metabolizing efficiencies (99 and 98%) have been reported so far for *Rhodococcus erythropolis* and *Alcaligenes xylosoxidans* respectively (40, 41). The isolates with minimal toluene removal were found to be IUBT23 (28% ± 4.24) and IUBT26 (50% ± 9.19). However, in literature minimum toluene removal efficiency has been reported to be 43% and 49% in rhizosphere bacterial community (42).

### Antibiotic profiling

All of the twenty bacteria reported can be used for toluene remediation. But there may be some associated potential risks due to their virulent nature. Although this study has not detected virulence status of isolates, the antibiotic resistance profiling may give us a clue about their user friendly nature. Environment friendly bacteria should not possess antibiotic resistance. Moreover, the exploitation of these bacteria to remediate toluene can be decided on the bases of their sensitivity as well as resistance to the drugs. Application of a whole bacterial cell to deal unfavorable toluene burden can only be suggested, if it is drug sensitive. Hence, it was crucial to test the isolated bacteria for antibiotic resistance. All of the bacteria were gram positive so we selected the antibiotics which are most commonly used for treatment of infections caused by gram positive bacteria. i.e; teicoplanin, linezolid, oxacillin and chloramphenicol. All of the bacteria in our study have shown resistance to some of the antibiotics so the whole cell remediation will not be recommended. Rather their enzymes and genes associated with toluene degrading pathways can be beneficially used and three approaches should be followed in this regard.

(a) If the required enzymes are extracellular then nanoparticles can be synthesized by using supernatant and can be destined for remediation.
(b) If intracellular enzymes are associated with toluene catabolism then these enzymes and other proteins can be purified from cell lysate for synthesis of nanoparticles.
(c) Associated genes can also be identified and can be cloned into any environmental friendly bacteria for bioremediation.

## Conclusion

Twenty thermophilic bacteria have been isolated which are capable of toluene catabolism. Biochemical analysis has revealed other catabolic attributes possessed by these bacteria which can be exploited for different industrial applications. There is still need to elucidate the pathways associated with toluene degradation. Study of genes encoding enzymes, factors as well as conditions contributing to maximum toluene breakdown would help to exploit their potential in an environmentally favorable way. By combining knowledge about these attributes of bacteria and bioremediation approach, toluene degrading potential of these bacteria can be harnessed to mitigate toluene pollution in the environment.

